# Emerging Accessibility Patterns in Long Telomeric Overhangs

**DOI:** 10.1101/2021.11.24.469879

**Authors:** Sajad Shiekh, Golam Mustafa, Sineth G. Kodikara, Mohammed Enamul Hoque, Eric Yokie, John J. Portman, Hamza Balci

**Affiliations:** Department of Physics, Kent State University, Kent, OH 44242, USA; Department of Chemistry and Biochemistry, Kent State University, Kent, OH 44242, USA

**Keywords:** Telomere, G-quadruplex, single molecule, FRET-PAINT

## Abstract

We present single molecule experimental and computational modeling studies investigating the accessibility of human telomeric overhangs of physiologically relevant lengths. We studied 25 different overhangs that contain 4-28 repeats of GGGTTA (G-Tract) sequence and accommodate 1-7 tandem G-quadruplex (GQ) structures. Using FRET-PAINT method, we probed the distribution of accessible sites via a short imager strand, which is complementary to a G-Tract and transiently binds to available sites. We report accessibility patterns that periodically change with overhang length and interpret these patterns in terms of the underlying folding landscape and folding frustration. Overhangs that have [4n]G-Tracts, (12, 16, 20…), demonstrate the broadest accessibility patterns where the PNA probe accesses G-Tracts throughout the overhang. On the other hand, constructs with [4n+2]G-Tracts, (14, 18, 22…), have narrower patterns where the neighborhood of the junction between single and double stranded telomere is most accessible. We interpret these results as the folding frustration being higher in [4n]G-Tract constructs compared to [4n+2]G-Tract constructs. We also developed a computational model that tests the consistency of different folding stabilities and cooperativities between neighboring GQs with the observed accessibility patterns. Our experimental and computational studies suggest the neighborhood of the junction between single and double stranded telomere is least stable and most accessible, which is significant as this is a potential site where the connection between POT1/TPP1 (bound to single stranded telomere) and other shelterin proteins (localized on double stranded telomere) is established.

**Significance Statement:** The ends of eukaryotic linear chromosomes are capped by telomeres which terminate with a single-stranded overhang. Telomeric overhangs fold into compact structures, called G-quadruplex, that inhibit access to these critical genomic sites. We report single molecule measurements and computational modeling studies probing the accessibility of a set of human telomeric overhangs that covers a significant portion of the physiologically relevant length scale. We observe novel accessibility patterns which have a well-defined periodicity and show that certain regions are significantly more accessible than others. These accessibility patterns also suggest the underlying folding frustration of G-quadruplexes depends on telomere length. These patterns have significant implications for regulating the access of DNA processing enzymes and DNA binding proteins that can target telomeric overhangs.

## Introduction

The ends of linear chromosomes, called telomeres, contain repeating sequences and play vital roles in promoting the integrity of these important genomic sites. Telomeres are involved in safeguarding the chromosome ends by differentiating them from DNA double-strand breaks, which would otherwise trigger unwanted DNA damage response (1, 2). Human telomeric DNA is composed of tandem repeats of hexanucleotide d(GGGTTA) arranged into a long double-stranded region followed by a 50-300 nucleotide (nt) long 3’ single-stranded overhang (3). These unique telomeric repeats are conserved across vertebrates and beyond (4). The length of telomeres is important for their protective functions (5, 6); however, the inability of DNA replication machinery to complete the replication and processing of chromosome ends causes progressive telomere shortening during successive cell divisions (7). When a critical telomere length is reached, apoptosis or cell senescence is activated in somatic cells (8). Certain oncogenic cells continue to proliferate by overexpressing telomerase ribonucleoprotein complex which elongates telomeres, eventually leading to development of malignant tumors (6, 9). Telomerase includes a reverse transcriptase enzyme that synthesizes telomeric DNA repeats d(GGTTAG) using an internal RNA template (3, 10).

The G-rich telomeric overhangs fold into compact non-canonical structures called G-quadruplex (GQ). GQ is formed when four repeats of d(GGGTTA) are arranged into planar quartet geometry, stabilized by Hoogsteen hydrogen bonding and centrally located monovalent cation. For brevity, the d(GGGTTA) sequence will be referred to as a “G-Tract”. Potentially GQ forming sequences are concentrated in certain regions of the human genome, including telomeres and promoters, and GQs have been confirmed to form in vitro (11, 12) and in vivo (13–15). GQs are implicated in crucial biological processes, such as transcription, recombination, and replication (16, 17). Telomeric GQs can inhibit telomerase from elongating telomeres (18). While much is known about telomeres, many fundamental questions remain about their structure, the nature of the interactions between folded GQs, and accessibility of different regions of the telomeric overhang.

Most studies on GQs have typically focused on understanding the structure and function of “single” DNA or RNA GQ molecules (19–21) and their interactions with DNA binding proteins (22–24) and helicases (25–28). However, the presence of many G-Tracts in human telomeric overhangs allows formation of 2-10 tandem GQs with varying separations. How these tandem GQs interact with each other, and their folding characteristics would clearly impact telomere accessibility. Prior studies on DNA constructs that form multiple telomeric GQs have typically investigated the thermodynamic, kinetic, or structural characteristics of isolated DNA molecules but have not directly studied telomere accessibility, which is the primary goal of this study. The studies on isolated tandem GQs have not reached a consensus and have either concluded negligible interactions, stabilizing stacking interactions (positive cooperativity), or destabilizing interactions (negative cooperativity) that take place between neighboring GQs (29–34). Variations in ionic conditions, temperature, sensitivity of probes to different thermodynamics parameters, or the design of the DNA constructs might have played a role in these different conclusions.

In this study, we investigated the accessibility patterns of telomeric single-stranded DNA (ssDNA) molecules that contain 4-28 G-Tracts, a total of 25 constructs, using the single molecule FRET-PAINT (Förster Resonance Energy Transfer-Point Accumulation for Imaging in Nanoscale Topography) method (34). These constructs can form 1-7 tandem GQs or could have multiple unfolded G-Tracts at different segments of the overhang. FRET-PAINT combines attributes of the super-resolution microscopy technique DNA-PAINT (35) with those of single-molecule FRET (smFRET) (36). This methodology takes advantage of transient binding of a labeled probe (called imager strand) to a target strand. As the imager strand, we used an acceptor (Cy5) labeled Peptide Nucleic Acid (PNA) that is complementary to a 7-nt segment of telomeric sequence. The imager strand is introduced to a microfluidic chamber that contains surface-immobilized, donor (Cy3) labeled partial duplex DNA (pdDNA) constructs that have a single stranded telomeric overhang (Figure 1A). This partial duplex design mimics the physiological structure. Binding of Cy5-PNA to an available G-Tract results in a FRET signal (Figure 1B), which is used as an indicator for telomere accessibility. This FRET signal depends on the position of the binding site within the overhang and the folding pattern between the binding site and the ssDNA/dsDNA junction, where Cy3 is located. The resulting FRET distributions represent the accessibility patterns of the overhang. The distinguishing feature of this novel approach is providing a direct probe of telomere accessibility for overhangs with physiologically relevant lengths.

**Figure 1.**
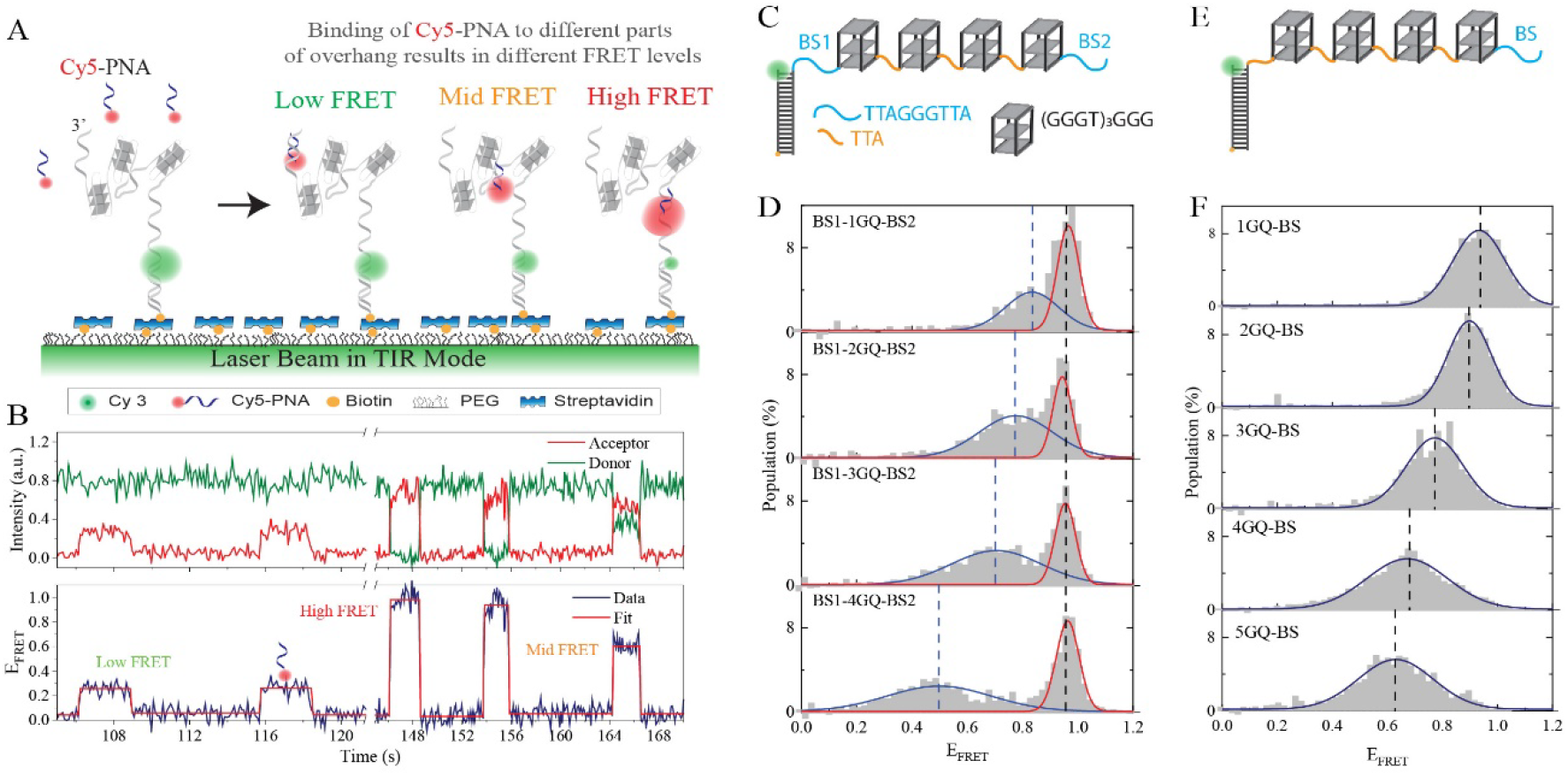
Schematic of FRET-PAINT assay. (A) Partial duplex DNA constructs (labeled with donor fluorophore Cy3) are immobilized on the PEGylated surface via biotin-streptavidin attachment. The pdDNA construct has a telomeric overhang with multiple GGGTTA (G-Tract) repeats, which can form GQs separated from each other with unfolded regions of varying length. Different FRET levels are observed when the imager strand (Cy5-PNA) transiently binds to different segments of the overhang. (B) An example smFRET time trace showing five binding events with low, mid, and high FRET levels. The red line is a fit to the FRET trace to indicate the corresponding FRET levels. (C) Schematics of overhangs that contain four GQs with a modified sequence [(GGGT)_3_GGG for each GQ] and two specific binding sites (BS1 and BS2) for Cy5-PNA. (D) FRET-PAINT data demonstrating the accessibility patterns for constructs in which BS1 and BS2 are separated from each other by 1-4 GQs. The FRET level representing binding to BS1 remains constant while that representing binding to BS2 gradually decreases but remains within detectable range. The blue and red curves are gaussian peak fits to the data (listed in Table S5). (E) Schematics of overhangs that contain four modified GQs and a specific binding site (BS) for Cy5-PNA. (F) FRET-PAINT data on constructs in which BS is separated from the junction by 1-5 modified GQs. The FRET level representing binding to BS systematically decreases as the overhang length increases yet it remains within detectable range even after 5-GQs. The blue curves are gaussian peak fits to the data (listed in Table S6), and dashed lines indicate the peak positions.

As accessibility patterns depend on the underlying GQ folding landscape, we also studied their implications for folding characteristics. For this, we developed a computational model that calculates the resulting accessibility patterns when folding stabilities for different segments of the overhang and nature of interactions between neighboring GQs are modulated. With this approach, we mapped the resulting patterns for a broad range of stability constants and investigated the resulting patterns for positive, negative and negligible cooperativity between neighboring GQs.

## Results

Before presenting the measurements on telomeric overhangs, we start with important control measurements that establish the validity and sensitivity of the FRET-PAINT approach for sequences that form tandem GQ structures. In these studies, we replaced the telomeric repeats (GGGTTA)_3_GGG with the (GGGT)_3_GGG sequence which forms a very stable GQ structure that attains only a parallel folding conformation (22). This modified sequence is complementary to only four consecutive nucleotides on Cy5-PNA, which is not long enough to form detectable PNA-DNA duplexes. To enable Cy5-PNA binding to specific locations, we inserted the TTAGGGTTA sequence as a binding site (BS) in one location (Figure 1E-F) or two locations (BS1 and BS2 in Figure 1C-D) on the overhang. In Figure 1C-D, the two binding sites were separated by 1-4 GQs, while in Figure 1E-F the binding site was separated from the junction by 1-5 GQs. To illustrate, the overhang shown in Figure 1C has the following sequence: TT<u>AGGGTTA</u> [(GGGT)_4_TA(GGGT)_4_TA(GGGT)_4_ TA(GGGT)_4_] T<u>AGGGTTA</u>. The sequence within the square brackets [ ] forms four GQs while the underlined sequences serve as the binding sites (BS1 and BS2 in Figure 1C) for Cy5-PNA.Complete sequences for these constructs are given in Table S1.

In the accessibility patterns of constructs with BS1 and BS2, we expect to have a high FRET peak that remains constant (representing binding to BS1) and a lower FRET peak (representing binding to BS2) that gradually shifts to lower levels as the overhang length increases. We also expect the lower FRET peak to broaden with length as the number of potential folding patterns increases, resulting in variability in detected FRET levels. The data in Figure 1D agrees with these expectations. The high FRET peak at E_FRET_≈0.95 remains unchanged in all constructs while the lower FRET peak gradually shifts to lower levels and broadens. Significantly, the FRET levels remain well within the detectable range even when BS2 is separated from the donor molecule by an unfolded G-Tract (BS1) and four GQs.

Figure 1F shows accessibility patterns for the constructs that have a single binding site separated from the junction by 1-5 GQs. To illustrate, the sequence of the overhang in Figure 1E is: TTA[(GGGT)_4_TA(GGGT)_4_TA(GGGT)_4_TA(GGGT)_4_] T<u>AGGGTTA</u>, where the underlined sequence serves as the binding site (BS in Figure 1E) for Cy5-PNA. As shown in Figure 1F, even after 5 GQs, the binding events to BS remain well within detectable FRET range. We conclude that the FRET-PAINT approach can be used to study the accessibility patterns of long telomeric sequences, which will be presented next.

Figure 2 shows the normalized FRET distributions obtained from transient bindings of Cy5-PNA to unfolded G-Tracts in telomeric overhangs. The data are grouped in sets of four histograms, with the shortest construct in each set having an integer multiple of 4 G-Tracts, [4n], while the longest construct has an additional three G-Tracts, [4n+3]. These FRET distributions reveal surprising accessibility patterns that emerge when the telomeric overhang reaches a length of about 10 G-Tracts. The patterns have a periodicity of 4 G-Tracts and persist in the 10-28 G-Tract range. While the [4n]G-Tract constructs have the broadest distributions, the [4n+2]G-Tract constructs are typically the narrowest. These narrower distributions are concentrated at high-FRET levels, which represent binding to sites closer to the ssDNA/dsDNA junction (5’-side). The broad distributions of [4n]G-Tract constructs suggest the binding sites, i.e., unfolded G-Tracts, are distributed throughout the overhang.

**Figure 2.**
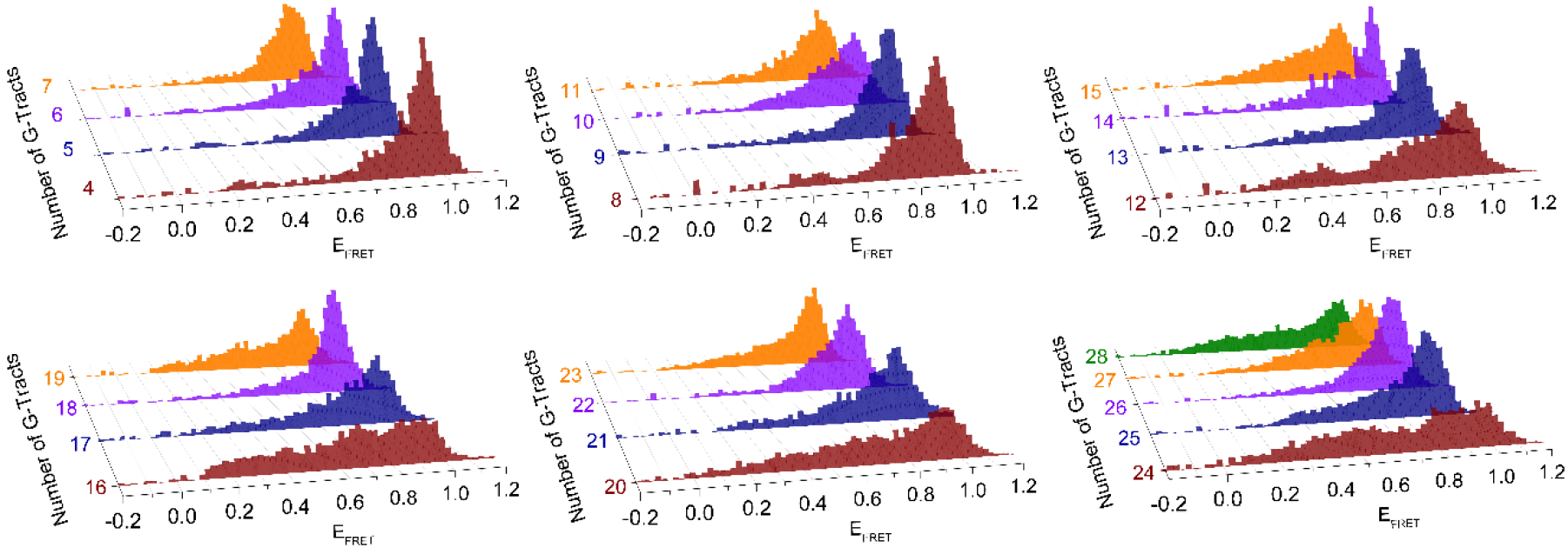
Normalized FRET histograms for telomeric overhangs with 4-28 G-Tracts. The histograms are grouped in sets of 4 with [4n]G-Tracts in brown, [4n+1]G-Tracts in blue, [4n+2]G-Tracts in violet and [4n+3]G-Tracts in orange. The 28G-Tract construct is added to the last histogram in green. The FRET histograms, which represent accessibility patterns, are broadest in [4n]G-Tract constructs and narrowest in [4n+2]G-Tract constructs.

Figure 3A shows a contour plot of the combined data for all 25 constructs that illustrates alternating broad (marked with white dashed lines) and narrow (marked with white dotted lines) distributions. To quantify the broadness of FRET distributions, we defined an S-parameter (Shannon entropy divided by the Boltzmann constant). The S-parameter describes the uncertainty associated with identifying the binding site of Cy5-PNA, which is greater for broader distributions. Therefore, the maxima of S-parameter occur at [4n]G-Tract constructs and the minima at [4n+2]G-Tract constructs (Figure 3B).

**Figure 3.**
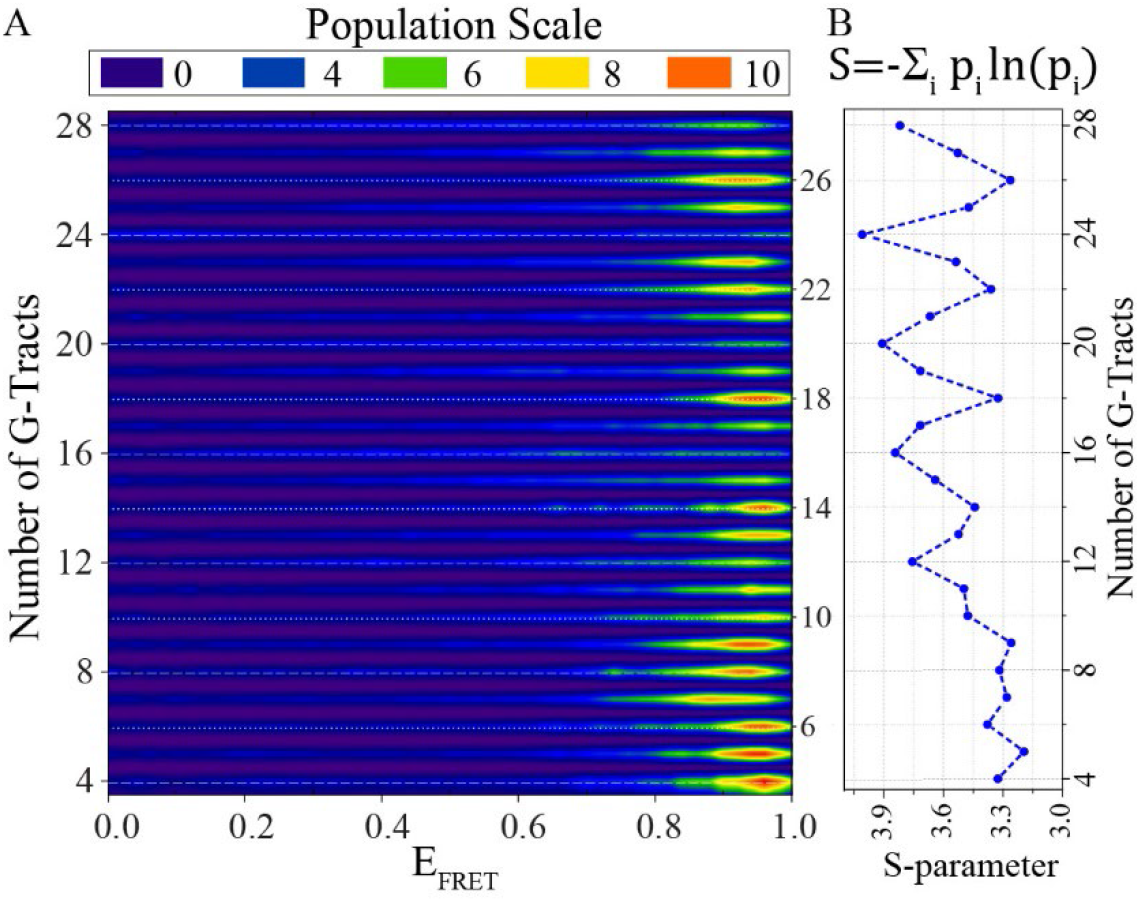
Contour plot of accessibility patterns and variation of S-parameter with overhang length. (A) The histograms in Figure 2 are combined in a contour plot that demonstrates broad histograms for [4n]G-Tract constructs (for 12G-Tract and longer) and narrower histograms for other constructs, especially [4n+2]G-Tract constructs. The overhangs with [4n]G-Tracts are marked with dashed white lines while [4n+2]G-Tract constructs are marked with dotted white lines. The same bin-size was used in this contour plot as that used in the histograms in Figure 2. (B) To quantify the broadness of the histograms in Figure 2, an S-parameter (defined on top) is introduced. S-parameter is a measure of folding frustration: the larger the S-parameter the greater the folding frustration. At lengths longer than 12G-Tract, a pattern is established where the maxima of S-parameter occur at [4n]G-Tract constructs while minima are at [4n+2]G-Tract constructs.

The observed accessibility patterns have implications about the underlying folding topography. We explored this connection through a computational model introduced by Mittermaier *et al*. (33). In this two state GQ model with nearest neighbor interactions (analogous to Ising model), combinatorial factors associated with the location of the folded GQs along the telomere and its finite length conspire to produce non-trivial accessibility patterns. This model predicts folding probability (P_i_) of a G-Tract based on folding cooperativity (K_c_) between neighboring GQs and the statistical weight of a folded GQ (K_f_). Due to reasons described in Materials and Methods section, the following parameters were adopted from Mittermaier *et al*. study: K_f_ = 60 for the stability of a GQ in the interior, K_f,N_ = 229 for a GQ ending at the 3’-end (for an overhang that includes N G-Tracts), and K_f,N-1_ = 80 for a GQ ending at the neighboring G-Tract. Our model predictions of folding patterns are robust as long as K_f,N-1_ is greater than K_f_ and K_f,N_ is significantly greater than K_f,N-1_. The G-Tracts on the 5’ side (vicinity of ssDNA/dsDNA junction) are significantly less stable due to the flanking duplex regions, similar to a recent observation in KIT promoter (37). We estimated the folding parameters for these 5’ G-Tracts (K_f,1_ and K_f,2_) and the cooperativity parameter (K_c_) by comparing the patterns observed for the experimentally measured and computationally calculated S-parameters, as described below.

The FRET signal measured in the experiment probes the location of unfolded G-Tracts. To compare the outcomes of the model with the experimental distribution measured as a function of telomere length, we calculate how delocalized the distribution of unfolded G-Tracts is for a given length via the S-parameter, as defined in the Materials and Methods section. We find that the model accommodates a variety of patterns of the calculated S-parameter as a function of telomere length depending on folding and cooperativity parameters used in the model. Figure 4 shows a phase diagram as a function of the stability for a GQ starting with the 5’ G-Tract, K_f,1_, and the cooperativity K_c_. In this figure, we choose the weight for a GQ starting at the second G-Tract from the junction to be twice that of starting at the first G-Tract, i.e. K_f,2_ = 2K_f,1_. Similar patterns are observed as long as K_f,2_ is greater than K_f,1_ and significantly smaller than K_f_ (Supplementary Figure S1). For simplicity, we focus on a window composed of N=20 to N=24 G-Tracts to characterize the pattern of the S-parameter and monitor if the S-parameter increases or decreases as N increases within this range. With positive cooperativity, four patterns are observed for the S-parameters. The pattern observed in the FRET efficiency (colored green in Figure 4) occurs within a specific (though rather broad) range for parameters K_f,1_, K_f,2_, and K_c_. For K_c_ >6, the stabilities of GQ starting near the ssDNA/dsDNA junction must be below an upper threshold to observe the appropriate pattern, and for smaller cooperativity, these stabilities must lie between a lower and upper bound. The S-parameter as a function of N, shown in Figure 4, agrees qualitatively with the measured pattern. Because of this similarity to the experimentally measured S-parameter, we will use parameters (K_f,1_ = 5, K_f,2_ = 10, and K_c_ = 5) as an illustration in what follows.

**Figure 4.**
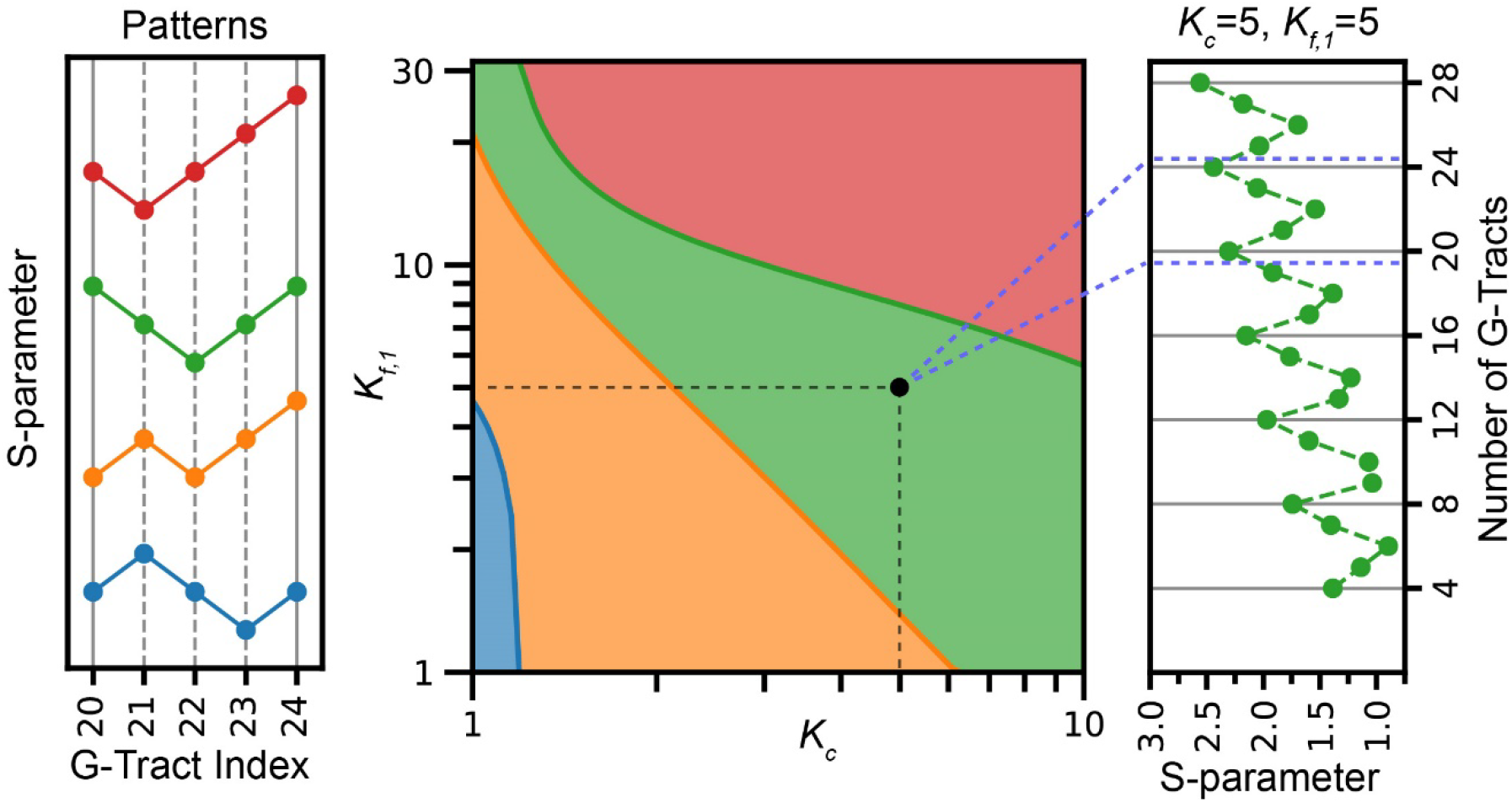
Computational patterns observed for the S-parameter for N=20-24 G-Tracts for varying K_f,1_ and K_c_ parameters. In these calculations, K_f,1_ and K_c_ are varied while K_f,2_=2K_f,1_, K_f_ =60, K_f,N-1_=80, and K_f,N_=229. The left panel shows the four patterns observed depending on whether the S-parameter increases or decreases as the length of the overhang is increased by one G-Tract. The phase diagram in the middle shows the parameter range where different patterns are observed in colors that match the patterns on the left panel. The green range corresponds to the pattern observed in the experiment. The right panel shows the calculated S-parameters for K_c_=5 and K_f,1_=5, which can be compared to the experimentally observed pattern in Figure 3B.

Figure 5 shows that the parameters identified by comparison of the computational and experimental S-parameters correspond to very stabilized GQs. This is consistent with the experiment where most of telomeric constructs do not show any binding events during experimental observation time (∼2-3 min). Although, the strong stability weakens as the number of G-Tracts increases, the average number of folded GQs (<n_GQ_>) is nearly maximal for G-Tracts ranging from N = 4 to 28 (Figure 5A). Nevertheless, constructs with 4n and 4n+1 G-Tracts have higher average number of unfolded G-Tracts (<n_uf_>) than the minimum associated with fully folded GQs (Figure 5B). Also, constructs with 4n and 4n+1 G-Tracts have large relative fluctuations, suggesting that the number of sites available to PNA binding can be significantly larger than indicated by <n_uf_> for these constructs.

**Figure 5.**
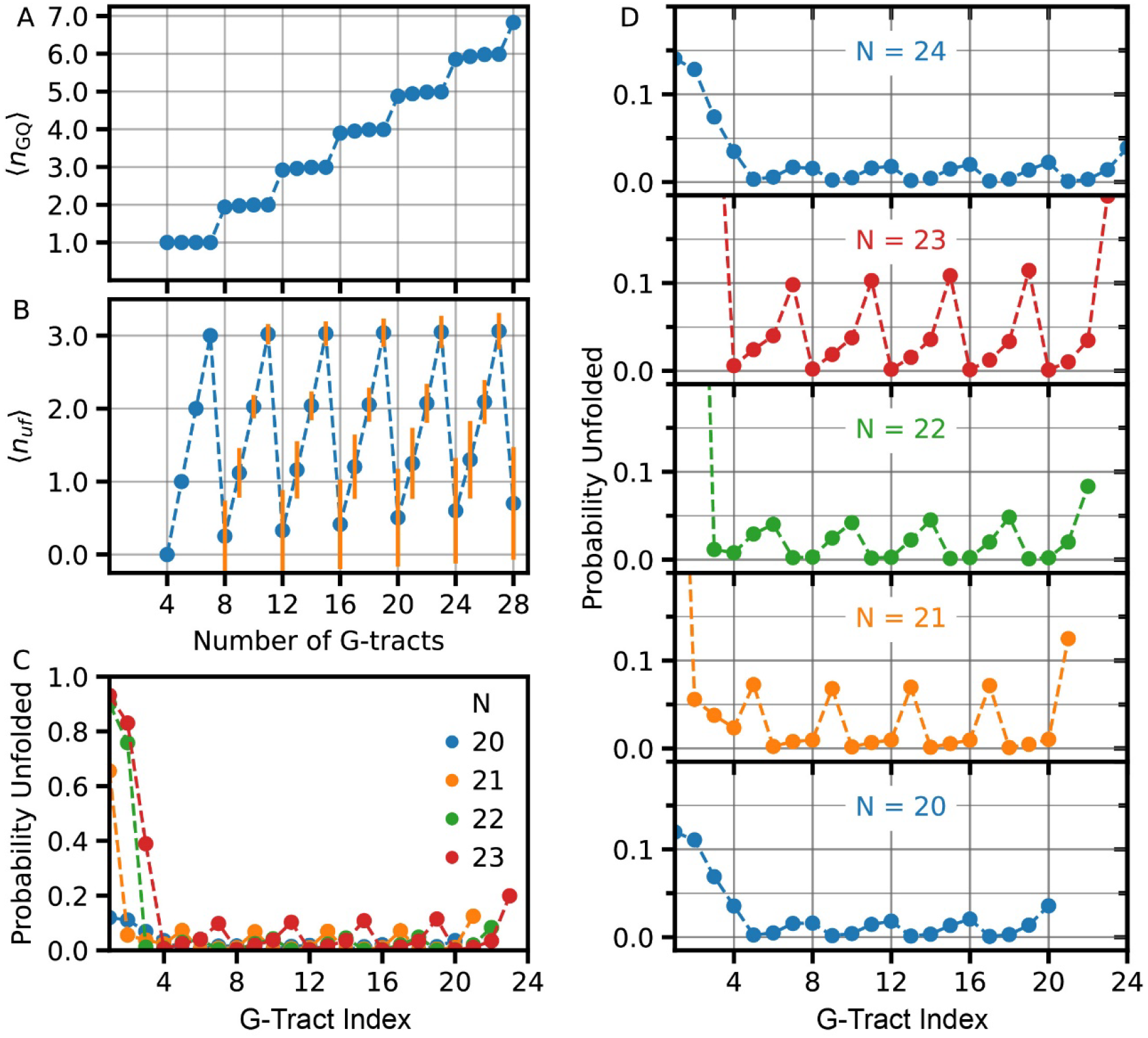
Folding levels for the set of parameters identified in Figure 4: K_f,1_=5, K_f,2_=10, K_f_ =60, K_f,N-1_=80, K_f,N_=229 and K_c_=5. These parameters result in high levels of folding for all constructs investigated although periodic patterns were observed for probability of being unfolded depending on the position of the G-Tract and length of overhang. (A) For the set of parameters used in these studies, the average number of GQ (<n_GQ_>) is close to the maximum possible throughout 4-28 G-Tract overhang length. (B) Average number of unfolded G-Tracts (<n_uf_>) as a function of overhang length. As expected, constructs with 4n+1, 4n+2, and 4n+3 G-Tracts have at least 1, 2, and 3 unfolded G-Tracts, respectively. As can be most clearly seen for 4n constructs, <n_uf_> increases as the overhang length increases. The orange vertical lines are standard deviation in <n_uf_>. (C) Probability of i^th^ G-Tract being unfolded as a function its position for overhangs with N=20-24 G-Tracts. In this diagram i=1 for the G-Tract at ssDNA/dsDNA junction and i=N for the G-Tract at 3’-end. (D) The plots that are overlaid in (C) are separated for each N, and the y-axis scale is limited to unfolding probability in 0.0-0.2 range (0.0<P_i_<0.2) to better illustrate the oscillations in the unfolding probability. The amplitude of oscillations in P_i_ varies depending on overhang length, which can be related to the patterns observed for S-parameter.

Patterns in the S-parameter can be understood as balancing the population of unfolded G-Tracts within the destabilized junction region, and the rest of the telomeric G-Tracts given the underlying frustration of the landscape and the constraint that all states have a minimum number of unfolded G-Tracts when the length of the construct differs from 4n. Because GQ involving the first two G-Tracts near the junction (at the 5’) side are destabilized compared to others, the unfolded distribution is shifted towards the first two repeats. As shown in Figure 5C-D, the G-Tracts are mostly folded for N = 4n, with a weak pattern in unfolding propensity along the chain. For N = 4n+1, which has at least one unfolded G-Tract, the first G-Tract is much more likely unfolded, and the pattern for the unfolded G-Tract along the chain is more prominent. This localization of the unfolded G-Tracts is reflected in a decrease in the S-parameter. For N = 4n+2, the first two G-Tracts are unfolded with high probability and the remaining unfolding propensity decreases along the chain. Thus, the distribution of Pi is more localized and results in a local minimum of the S-parameter. For N = 4n + 3, unfolding of the first three G-Tracts are more likely, though the propensity of unfolding for other G-Tracts increases, giving rise to a more delocalized distribution along the chain, and hence an increase in the S-parameter. A similar analysis of the patterns of the S-parameters shows that the pattern observed in the experiment does not develop when folding cooperativity is negative (Supplementary Figure S2). Also, looking at a broader window N = 12 – 24, we see that the pattern can shift from one type to another as a function of N near the boundaries of phases of the S-parameter (Supplementary Figure S3). Furthermore, these kinds of patterns occur not only in the S-parameter which explicitly depends on the distribution of unfolded G-Tracts, but also in the entropy as a function of N (Supplementary Figure S4).

## Discussion

This study differs in significant aspects from most studies in literature and provides important findings that were not accessible previously. Firstly, we employ DNA constructs that contain a duplex DNA on one side (similar to the physiological telomeric DNA) rather than two free ends as done in most studies. Secondly, we directly probe accessibility of the telomeric overhang to a complementary PNA probe and interpret the results in terms of their implications for relative stability of different segments of the overhang and interactions between neighboring GQs. Most studies in literature follow the opposite route where telomeric DNA is studied in isolation and stabilities of different segments are determined. These stabilities are then interpreted in terms of their implications for the accessibility of the overhang. To our knowledge, the accessibility maps demonstrated in Figure 3 are the first demonstration of this kind of a direct approach.

The PNA probe we employ in our studies creates a small disturbance to the underlying folding topography. This approach has physiological relevance as is evident from the cases of telomerase or TERRA, which contain nucleic acids templates that interact with telomeres and impact the underlying GQ structures. Our Cy5-PNA probe base-pairs with 7 nt telomeric overhang (TTAGGGT), which is smaller than the 9-nt long minimum unfolded site (TTAGGGTTA). To determine the thermodynamic stability of the PNA/DNA duplex that would form upon binding of Cy5-PNA to a telomeric sequence, we performed FRET melting measurements under identical ionic conditions to those of smFRET measurements and obtained a melting temperature of T_m_=12.7±0.3 °C (Supplementary Figure S5). This is consistent with the estimate of a model (T_m_=13.7 °C) that predicts T_m_ of a PNA/DNA duplex based on the melting point of a DNA/DNA duplex of the same sequence (38). Compared to the stability of a telomeric GQ (T_m_=68 °C) (39), this is a very weak disturbance. This is also evident in the low frequency of the binding events we observe and the fact that majority of DNA molecules that pass the single molecule screening test do not show any binding event during the experimental observation time (2-3 minutes) (Table S4). To provide a quantitative comparison, the frequency of binding events to a single G-Tract (an exposed G-Tract in the absence of GQ) is over an order of magnitude greater than the frequency of binding events for the 4G-Tract construct (Supplementary Figure S6). One might expect that there should not be any binding events for a 4G-Tract construct since all G-Tracts would be protected within a GQ. However, this assumes all the DNA molecules are perfectly folded and remain so throughout the observation time. Another piece of evidence for the Cy5-PNA probe not destabilizing the GQs in KCl comes from the observation that the frequency of binding events are much higher in LiCl compared to KCl, as we demonstrated in an earlier study (34).

The folding patterns of Figure 2 could be interpreted in terms of folding frustration where higher frustration would be expected to result in broader distributions. With this identification in mind, our data suggest [4n]G-Tract constructs have significantly higher frustration compared to other constructs. Adding 1-3 G-Tracts resolves this frustration to different extents, resulting in a concentration of unfolded sites in the vicinity of the ssDNA/dsDNA junction, on the 5’-side. The [4n]G-Tract constructs have an exact number of G-Tracts that can accommodate n GQs. Therefore, attaining complete folding requires a perfect progression of the folding throughout the overhang. For these constructs, nucleation of GQ folding from multiple sites will likely result in leaving unfolded G-Tracts between neighboring GQs. Also, skipping of even one G-Tract during progression of the folding (e.g., two neighboring GQs separated by an unfolded G-Tract) would result in three more unfolded G-Tracts at different regions of the overhang. We assume GQs with long loops that incorporate one or more G-Tracts (9 nt or longer loops instead of the canonical 3 nt loops) are highly unlikely due to entropic considerations (33). On the other hand, constructs with additional G-Tracts ([4n+1], [4n+2] and [4n+3]) are more tolerant for shifts in the G-Tract register during folding or initiation of folding from different sites.

Our data suggest the G-Tracts at the 3’-end or intermediate regions have higher folding stability compared to those at the 5’-side, which might be due to proximity of these sites to a duplex DNA. Therefore, nucleation of folding from the 3’-end or intermediate regions would more likely result in unfolded G-Tracts at the 5’-side. Once a particular folding conformation is established, it persists for long periods of time due to the high stability of associated GQ structures, kinetically hindering resolution of the frustration in folding (33). These observations have physiological significance as the free 3’-ends of telomeres are more prone to degradation by exonuclease activity and folding into GQ helps in protecting these ends (40). In addition, both telomerase and alternative lengthening of telomere (ALT) mechanisms require unfolded 3’-ends for the extension of telomeres (41) and presence of GQs could potentially render telomeres inaccessible. In addition, the ssDNA/dsDNA junction region, which is more likely to be unfolded, is a potential site where POT1-TPP1 might connect to other shelterin proteins, which are localized on duplex DNA (42). Therefore, having unfolded G-Tracts at this region would facilitate binding of POT1 and establishing this connection, which would also reduce accessibility of these regions to other DNA processing enzymes.

The conclusions of this study rely on the validity of FRET-PAINT approach to detect binding events throughout the long telomeric overhangs. For unstructured ssDNA, it would not have been possible to detect binding events 100 nt away from a reference site (location of donor fluorophore) using Cy3/Cy5 as FRET pairs. However, the telomeric overhangs are highly folded and stable (most DNA molecules do not show any binding events during the 2-3 min observation time) under our assay conditions. Therefore, the overhangs are much more compact than unstructured DNA. These considerations were supported with the experiments on modified constructs in Figure 1C-F. However, despite their low levels of abundance, the unfolded G-Tracts are physiologically very significant as they might serve as the nucleation sites for triggering DNA damage response, nuclease activity, or telomerase-mediated telomere extension.

Along these lines, it is also important to ensure that the compact form and 3D structure of the telomeric overhang do not eliminate the correlation between the observed FRET levels and telomere length. Potential interactions between neighboring GQs and the flexible segments (e.g., the TTA sequences that link consecutive GQs) in the telomeric overhang could impact this relationship. However, our data show that the correlation between the observed FRET efficiencies and telomere length are maintained. This is particularly evident in the histograms for [4n]G-Tract constructs where the binding sites are distributed throughout the overhang and hence provide more detailed information about the 3D structure compared to constructs where binding sites are concentrated in a particular region. In the [4n]G-Tract constructs, the FRET distributions systematically shift to lower FRET levels as the overhang length is increased, suggesting the correlation between FRET efficiency and location of binding site (with respect to the junction where the donor fluorophore is located) is not eliminated. We quantified this by studying the fraction of FRET population for E_FRET_<0.50 and demonstrate that this population increases as telomere length is increased (Supplementary Figure S7) (34). We note that, depending on the folding pattern, binding to different sites might result in similar FRET efficiencies or binding to a particular site could result in different FRET values in our FRET PAINT measurements. Therefore, we based our conclusions on the patterns and periodicities observed in the FRET distributions (Figure 3) rather than exact identification of binding sites. Also, we limited our statements to distinguishing three regions in the overhang: the junction region, the broad intermediate region, and the vicinity of the 3’-end.

Whether accessibility of a particular open site is impacted by its neighborhood is another relevant issue. To illustrate, an open site (a G-Tract that is not part of a GQ) could have folded GQs on both sides of it, a folded GQ on one side and another open site on the other, or open sites on both sides of it. Possible arrangements multiply if beyond nearest neighbors are considered. Since we do not have any control on which of these arrangements prevails for a given open site, we did not quantify the impact of the neighborhood structures on accessibility. However, since over a hundred binding events are used to create each of the histograms in Figure 2, these effects are expected to average out. Attaining more precise information would require performing systematic measurements on many constructs where certain configurations are biased by introducing mutations in the sequence, which will be the focus of future studies.

While the energy landscape of DNA GQ folding has been studied less extensively than that of proteins, it appears that the folding landscape of human telomeric GQ is not funneled (43) and has a high degree of frustration as evidenced by stable folding intermediates (44), alternative conformers (45), and sensitivity of the GQ topology to perturbations such as small changes in loop length (46). In this paper, the frustration does not refer to the to the frustration in folding of a single GQ (which we take as two-state in the model) but rather to the frustration associated with the folding patterns along the telomere in the sense of the term introduced by Mittermaier et al. who introduced the term frustration to describe the diversity of folding patterns in states with similar energy (33). This entropic and topological frustration has thermodynamic and kinetic consequences for accessibility of telomeres. In the present paper, we find that this simple model supports a surprising diversity of thermodynamic accessibility patterns which are sensitive to GQ interaction strength, stability, and telomere length. More broadly, frustration in biomolecules (47) is perhaps more familiar as a component of the funneled energy landscape of proteins (48). While natural protein sequences are minimally frustrated, landscape ruggedness and frustration is evident in patterns of frustration in many proteins (49), with locally frustrated regions often associated with allosteric conformation change (50) and protein-ligand binding sites (49). The frustration associated with folding of repeat proteins with coupled neighboring repeat units is perhaps most closely related to the frustration of GQ folding in telomeres. One-dimensional Ising models similar to the GQ model employed in this paper have been used to understand the folding thermodynamics and kinetics of repeat proteins (51, 52). In repeat proteins, the balance between the free energy of folding each element and the interactions between them is subtle, leading to a rich variety of behaviors. Consequently, proteins poised at a particular balance may be susceptible to local perturbations that cause large structural effects (51). The sensitivity of the accessibility patterns on the number of repeats (modulo 4) may be an analogous illustration of frustration in the telomeric landscape.

To get a feel for the stabilities suggested by this model, at T = 25 C, the stability of an interior GQ relative to an unfolded G-Tract is given by ΔG = -2.43 kcal/mol and a GQ involving the first G-Tract at the ssDNA/dsDNA junction is ΔG = -0.95 kcal/mol. The free energy of two adjacent GQ has free energy ΔG = -5.81 kcal/mol, of which -0.95 kcal/mol is associated with cooperative stabilizing interactions between the GQ’s. Destabilization of GQs at the ssDNA/dsDNA junction and positive cooperativity are consistent with the high FRET peak observed in many constructs. The free energies for GQ stability calculated from our model are consistent with those reported in literature (53); although, there is no consensus about whether the interaction between neighboring GQs is stabilizing or destabilizing. Several studies have investigated the interactions between neighboring GQs and possible higher-order structures that might form in telomeric overhangs using different constructs and experimental methods. However, these studies have not reached a consensus with conclusions varying from negligible stacking interactions (beads on a string type arrangement) (29) and destabilizing interactions (negative cooperativity) (33, 54) to higher order structures with multiple GQs condensing into compact structures (30, 34, 55, 56), which would suggest positive cooperativity. Our results are more consistent with these latter studies that propose positive cooperativity.

In this study, we demonstrate a direct probe of the accessible sites within telomeric overhangs of physiologically relevant lengths. This was made possible by the sensitivity of the single molecule methods employed in this study to minority populations, as these sites are only a small fraction of all potential sites on the telomeric overhangs. Our results demonstrate repeating accessibility and folding frustration patterns in these constructs. Overhangs with [4n]G-Tracts demonstrate elevated levels of frustration where the PNA probe is able to access sites throughout the overhang. On the other hand, overhangs with two additional G-Tracts, [4n+2]G-Tracts, show minimal frustration where most accessible sites are concentrated in the vicinity of ssDNA/dsDNA junction. Our computational studies also capture the requirement for lower stability at the ssDNA/dsDNA junction to attain patterns consistent with those observed in the experiment. These accessibility patterns would help with protecting the free 3’-end against exonuclease activity and inhibit telomerase-mediated elongation while facilitating binding of POT1/TPP1 in the vicinity of ssDNA/dsDNA junction and connect with other shelterin proteins.

## Materials and Methods

### Nucleic Acid Constructs

DNA strands were purchased either from Eurofins Genomics or IDT DNA and their sequences are given in Tables S1, S2 and S3. The oligonucleotides were purified in-house using denaturing polyacrylamide gel electrophoresis (PAGE). The corresponding gel images are shown in Supplementary Figure S8. HPLC purified Cy5-PNA strand was purchased from PNA-Bio Inc. The sequence of the Cy5-PNA probe is <u>TAACCCT</u>T-Cy5; the underlined nucleotides are complementary to 7-nt of telomeric sequence. The pdDNA constructs (Figure 1A) were formed by annealing a long strand that contains the telomeric overhang (4-28 G-Tracts) with a short strand (18, 24, or 30 nt) that has a biotin at the 3’-end and Cy3 at the 5’-end. The two strands were annealed in a thermal cycler in 150 mM KCl and 10 mM MgCl_2_ at 95°C for 3 minutes followed by a slow cooling to 30 °C (1 °C decrease every 3 minutes). This slow cooling process ensures attaining a thermodynamic steady state for the folding pattern. It is important to note that 10 mM MgCl_2_ was used only during the annealing process (to improve hybridization), while all measurements were performed at 2 mM MgCl_2_. During annealing, excess long strand (500 nM for 100 nM of short strand) was used to ensure all biotinylated and Cy3-labeled short strands have a matching long strand. The unpaired long strands are washed out of channel after the DNA is immobilized on surface via biotin-streptavidin linker.

To minimize its impact, the Cy5 is attached to PNA via a flexible linker and an additional thymine that does not hybridize with telomeric DNA. If Cy5-PNA binds in the immediate vicinity of a GQ, there will be 3 nt (TT from DNA and T from PNA) and 6-carbon long flexible linker that are not part of the GQ or the PNA/DNA duplex. Similarly, the Cy3 fluorophore at the junction (placed on the short strand) is separated by the nearest GQ by at least three nucleotides (TTA in the junction and 6-carbon long flexible linker). Earlier studies have shown that Cy3/Cy5 fluorophores placed at the terminal nucleotide tend to stack on the dsDNA (57). With 3-nt separation, Cy3 is not expected to have a significant impact on the nearest GQ. In most studies that employ fluorescent dyes (including bulk FRET melting assays), the fluorophores are separated from the GQ by 0-2 nt overhangs and a flexible linker. Even though these would not warrant zero interaction between the fluorophore and the GQ (58), the impact should be small and should be considered as an overall impact of the probe on the telomeric overhang.

### The smFRET assay and Imaging Setup

A home-built prism-type total internal reflection fluorescence (TIRF) microscope was used for these measurements following protocols described in earlier work (59). The slides and coverslips were initially cleaned with 1M potassium hydroxide (KOH) and acetone, followed by piranha etching, surface functionalization by amino silane and surface passivation by polyethylene glycol (PEG). To reduce nonspecific binding, the surfaces were treated with a mixture of m-PEG-5kDa and biotin-PEG-5kDa in the ratio 40:1 followed by another round of passivation with 333 Da PEG to increase the density of the PEG brush. The microfluidic chamber was created by sandwiching a PEGylated slide and a coverslip with double sided tape, followed by sealing the chamber with epoxy. The chamber was treated with 2% (v/v) Tween-20 to further reduce non-specific binding.

After washing the excess detergent from the chamber, 0.01 mg/mL streptavidin was incubated in the chamber for 2 minutes. The pdDNA samples were diluted to 10 pM in a buffer containing 150 mM KCl and 2 mM MgCl_2_ and incubated in the chamber for 2-5 minutes resulting in surface density of ∼300 molecules per imaging area (∼50 μm × 100 μm). The excess DNA was removed from the chamber with a buffer containing 150 mM KCl and 2 mM MgCl_2_. Unless otherwise specified, the imaging buffer contained 50 mM Tris-HCl (pH 7.5), 2 mM Trolox, 0.8 mg/mL glucose, 0.1 mg/mL glucose oxidase, 0.1 mg/mL bovine serum albumin (BSA), 2 mM MgCl_2_, 150 mM KCl and 40 nM Cy5-PNA. The Cy5-PNA strand was heated to 85 °C for 10 minutes prior to adding it to the imaging buffer. 1500-2000 frame long movies with 100 ms frame integration time were recorded using an Andor Ixon EMCCD camera. A green laser beam (532 nm) was used to excite the donor fluorophore and the fluorescence signal was collected by an Olympus water objective (60x, 1.20 NA).

### Data Analysis

A custom written C++ software was used to record and analyze the movies, and to generate single molecule time traces of donor and acceptor intensities. The time traces of each molecule were further analyzed by a custom MATLAB code to select the molecules that passed a single molecule screening test. The background was subtracted for each of these molecules based on the remnant donor and acceptor intensities after donor photobleaches. The FRET efficiency (E_FRET_) was calculated using E_FRET_ = Acceptor Intensity/(Acceptor Intensity + Donor Intensity). The molecules that did not show any binding event formed a donor-only (DO) peak at E_FRET_ =0.06, which was used as a reference to rescale the FRET distribution such that DO peak corresponds to E_FRET_ =0.00. After this rescaling, the DO peak was subtracted from the histograms. Therefore, the FRET histograms presented in Figure 2 were generated from single molecules that show at least one Cy5-PNA binding event. The numbers of molecules that contributed to each histogram in Figure 2 are presented in Table S4, but typically each histogram was constructed from binding events to 100-150 DNA molecules, resulting in several hundred binding events in each histogram. The constructs that do not have any unfolded G-Tracts or those whose unfolded sites are not accessible to the probe do not contribute to the histograms in Figure 2.

The FRET distributions were normalized such that each molecule contributes equally to the histogram and the total population across the entire FRET range was normalized to 100%, i.e., the population of a particular FRET bin represents it’s percent population. The probability *p*_*i*_ of a particular FRET level _*i*_ (a FRET bin of width 0.02) is obtained by dividing population at that FRET level by 100. To quantify the spread in the FRET histograms, we define an S-parameter: *s* = − ∑_*i*_ *p*_*i*_ In *p*_*i*_ where the summation is carried over the entire FRET range and *p*_*i*_ is the unfolded probability, normalized so that 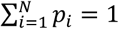. These calculations were performed in Origin 2015.

### Computational Model

We consider a simple model introduced by Mittermaier and co-workers to analyze the folding patterns in a telomeric sequence consisting of N G-Tracts, each of which can either be unfolded or folded with three neighboring G-Tracts in a GQ (33). The G-Tracts are counted from the junction (*i* = 1) towards the 3’-end (*i* = *N*). Mittermaier *et al*. used ssDNA constructs with two free ends while we used pdDNA constructs in which the telomeric overhang has a free 3’-end but borders a double stranded DNA on the 5’-side, similar to the physiological telomeres. Due to the similarity of our pdDNA constructs to the ssDNA constructs of Mittermaier *et al*. study in the middle and 3’-ends, we kept the stabilities of a GQ in the interior (K_f_ = 60), a GQ ending at the 3’-end (K_f,N_ = 229, for an overhang that includes N G-Tracts), and a GQ ending at the neighboring G-Tract (K_f,N-1_ = 80) similar to those reported in Mittermaier *et al*. study (33). On the 5’ side, the stability of the G-Tracts in our pdDNA constructs are significantly lower than those in ssDNA constructs of Mittermaier *et al*. study as we have a flanking duplex region, rather than a free end.

The statistical weight for a state α with 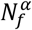 folded GQs and 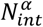 interfaces between nearest neighbor GQs is given by 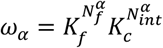, where *K*_*f*_ is the weight associated with a folded GQ, and *K*_*c*_ models interactions between adjacent GQs. Folding cooperativity between neighboring GQs can be positive (*K*_*c*_ > 1), negative (*K*_*c*_ < 1), or neutral (*K*_*c*_ *=* 1). In Mittermaier’s study, the DNA constructs were in the form of single stranded DNA with symmetrical 3’ and 5’ ends. This study demonstrated higher folding stability for GQs beginning or ending on the two terminal G-Tracts on each end: initiating on the 1^st^ and 2^nd^ G-tract at the 5’-end, or ending on the (N-1)^th^ and N^th^ G-Tract at the 3’-end. Compared to the internal G-Tracts: *K*_*f*,1_, *K*_*f*,2_, *K*_*f,N*−1_ and *K*_*f,N*_ were greater than *K*_*f*_ for internal G-Tracts. Since the free 3’-end and interior regions of our pdDNA constructs are identical to those of the Mittermaier study, we used similar statistical weights for *K*_*f*_, *K*_*f,N*−1_ and *K*_*f,N*_ as those estimated in that study. However, the vicinity of the dsDNA/ssDNA junction was significantly different from the rest in our constructs, which required using much lower folding parameters. These considerations are implemented as *K*_*f,N*_ > *K*_*f,N*−1_ > *K*_*f*_ > *K*_*f*,2_ > *K*_*f*,1_ in our study.

The probability for a state *α* is given by *p*_*α*_ = *ω*_*α*_ /*Z* where *Z* = ∑_*α*_ *ω*_*α*_ is the partition function. The total number of states can be written as 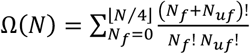, where the combinatorial factor is the number of states with *N*_*f*_ folded GQs and *Nu*_*f*_ = *N* − 4*N*_*f*_ unfolded G-Tracts. For telomeric overhangs of interest, Ω(*N*) is relatively small. For example, a telomere of 32 G-Tracts has only 16493 possible states, assuming long loops that contain one or more G-Tracts (such loops would be 9, 15, or 21 nt long compared to the canonical 3 nt loops) are not allowed due to their lower stability (33). This allows the partition sum to be computed explicitly by generating all states and evaluating the statistical weight of each state in the sum. The probability that the i^th^ G-Tract is unfolded is given by *P*_*i*_ *=* ∑_*α*_ *δ*_*i*_,_*α*_ *p*_*α*_, where *δ*_*i*_,_*α*_ = 1 if the ith G-Tract is unfolded and *δ*_*i, α*_ *=* 0 if the G-Tract is in a folded GQ. Overall, average properties of the telomere can be computed, such as the average number of folded GQs, 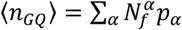, the average number of unfolded G-Tracts, ⟨*nu*_*f*_⟩ = *N* − 4⟨*n*_*GQ*_⟩, and the fluctuations of the unfolded G-Tracts, 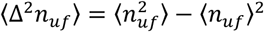.

For short chains, this model predicts a non-uniform unfolding probability along the sequence of G-Tracts under conditions strongly favoring folding (33). Here, combinatorial factors associated different ways in which specific G-Tracts can participate in GQ formation, as well as competing states with similar free energy conspire to produce patterns in *P*_*i*_ as a function of G-Tract index i; that is, some G-Tracts are more likely to be unfolded than others depending on their position. The experimental FRET distribution reflects binding of PNA to unfolded G-Tracts modeled by the distribution *P*_*i*_ which depends on the location of the G-Tract along the chain and the total number of repeats. This degree of localization of the unfolded G-Tracts is characterized by computing the S-parameter: 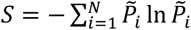, where 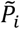 is the unfolded probability normalized to 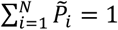

## Acknowledgments

This work was supported by NIH (1R15GM123443 to H.B.). We thank Dr. Soumitra Basu for use of his lab to perform the PAGE experiments.

