## Supplementary Information for "Emerging Accessibility Patterns in Long Telomeric Overhangs"

### Supplementary Information
### Emerging Accessibility Patterns in Long Telomeric Overhangs

Sajad Shiekh[a], Golam Mustafa[a], Sineth G. Kodikara[a], Mohammed Enamul Hoque[b], Eric Yokie[a], John J. Portman[a,*] and Hamza Balci[a,*]

[a] Department of Physics, Kent State University, Kent, OH 44242, USA;

[b] Department of Chemistry and Biochemistry, Kent State University, Kent, OH 44242, USA;

* To whom correspondence should be addressed:

**This PDF file includes:**

Supplementary text
Figures S1 to S8
Tables S1 to S6

**Computational Studies where $K_{f1}$ and $K_{f2}$ are Varied**

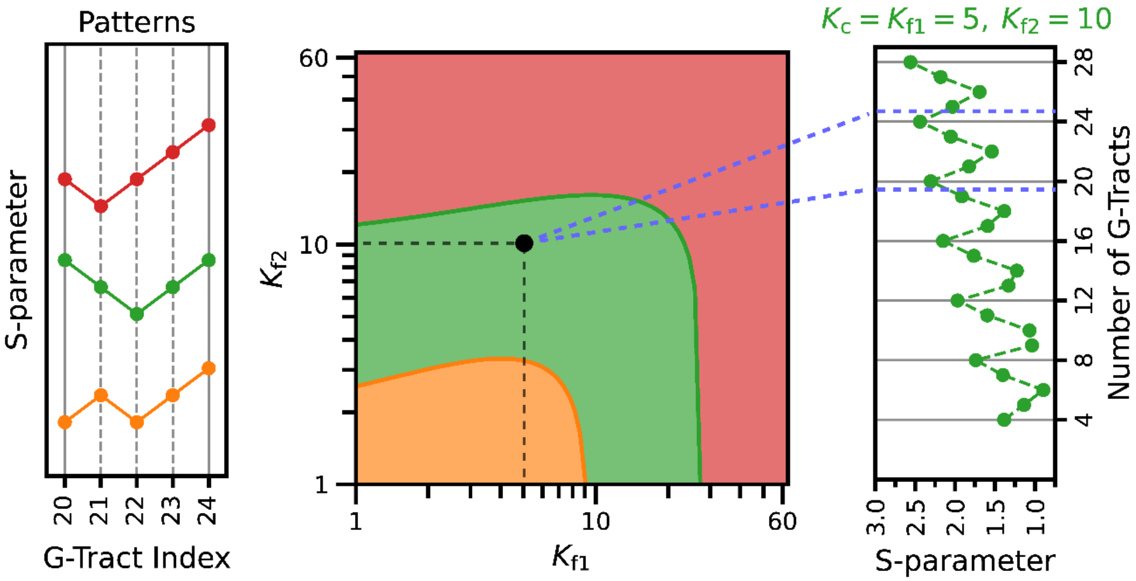


**Figure S1.** Computational patterns observed for the S-parameter for N=20-24 G-Tracts for varying $K_{f,1}$ and $K_{f,2}$ parameters. In these calculations, $K_{f,1}$ and $K_{f,2}$ are varied while $K_c$=5, $K_f$ =60, $K_{f,N-1}$=80, and $K_{f,N}$=229. The left panel shows the three patterns observed depending on whether the S-parameter increases or decreases as the length of the overhang is increased by one G-Tract. The phase diagram in the middle shows the parameter range where different patterns are observed in colors that match the patterns on the left panel. The green range corresponds to the pattern observed in the experiment. The right panel shows the calculated S-parameters for $K_c$=5, $K_{f,1}$=5, and $K_{f,2}$=10, which can be compared to the experimentally observed pattern in Figure 3B.

**Probing Negative Cooperativity**

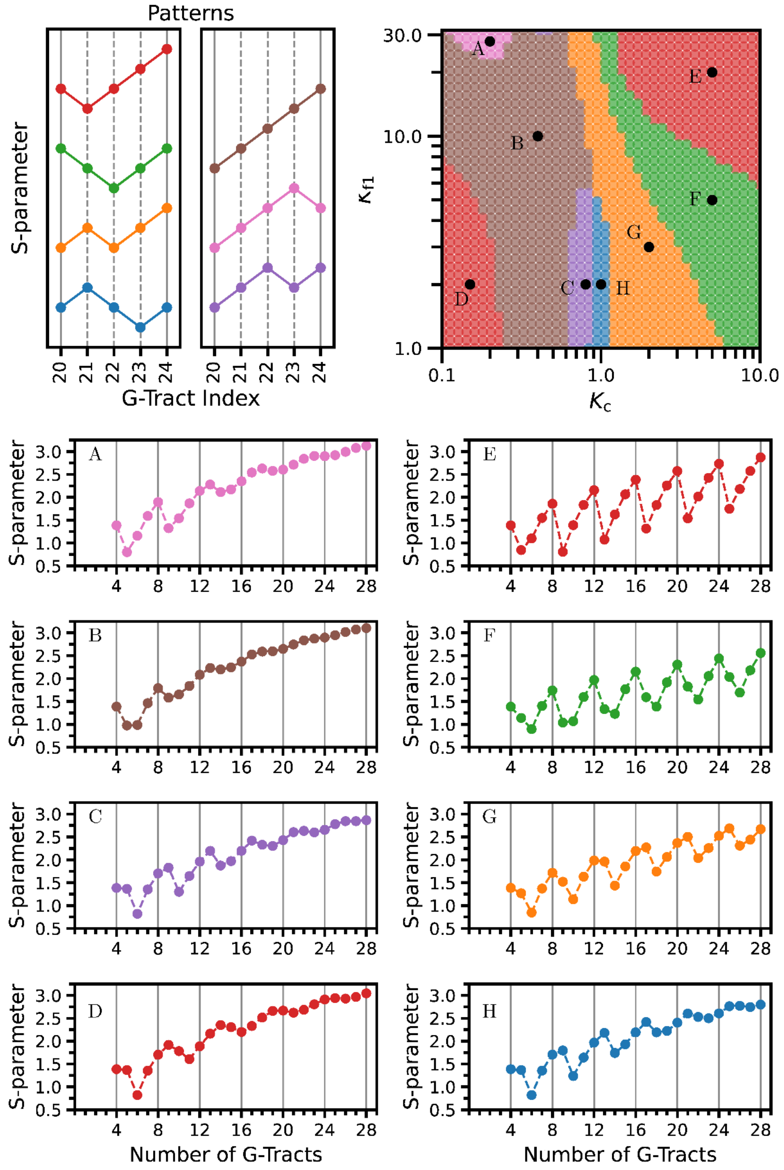


**Figure S2.** Computational patterns observed for the S-parameter for N=20-24 G-Tracts for varying $K_{f,1}$ and $K_c$ parameters. This plot expands the range of $K_c$ shown in Figure 4 to include negative cooperativity ($K_c$<1). In these calculations, $K_{f,1}$ and $K_c$ are varied while $K_{f,2}$=2$K_{f,1}$, $K_f$ =60, $K_{f,N-1}$=80, and $K_{f,N}$=229. The top left panel shows the seven patterns observed depending on whether the S-parameter increases or decreases as the length of the overhang is increased by one G-Tract. The phase diagram in the top right shows the parameter range where different patterns are observed in colors that match the patterns on the left panel. The S-parameters as a function of the number of G-tracts is shown in parts A – H. The parameters for the plots in A-H are labeled on the phase diagram. A: $K_c$ = 0.2, $K_{f,1}$ = 28; B: $K_c$ = 0.4, $K_{f,1}$ = 10; C: $K_c$ = 0.8, $K_{f,1}$ = 2; D: $K_c$ = 0.15, $K_{f,1}$ = 2; E: $K_c$ = 0.5, $K_{f,1}$ = 20; F: $K_c$ = 5.0, $K_{f,1}$ = 5.0; F: $K_c$ = 5.0, $K_{f,1}$ = 5.0; G: $K_c$ = 2.0, $K_{f,1}$ = 3.0; G: $K_c$ = 1.0, $K_{f,1}$ = 2.0. The green range (F) corresponds to the pattern observed in the experiment shown in Figure 3B. Notice that for negative cooperativity, the patterns in the S-parameters are prominent for small telomeres, but are significantly weaker for longer telomeres, greater than about N = 20.

**S-Parameter Patterns over a Broader Overhang Lengths**

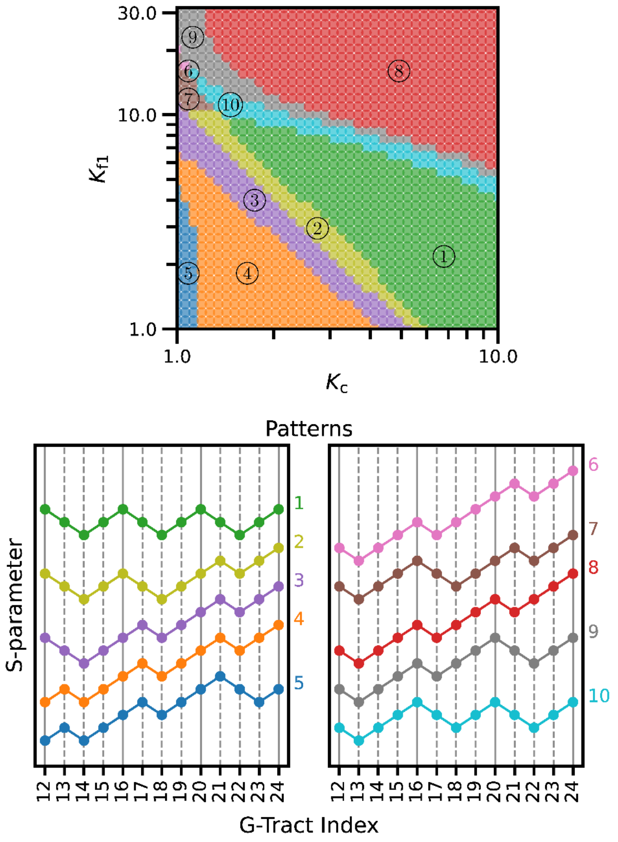


**Figure S3.** Computational patterns observed for the S-parameter for N=12-24 G-Tracts for varying $K_{f,1}$ and $K_c$ parameters. This calculation expands the window of G-Tracts used to classify patterns in the S-parameter in Figure 4, which uses only N=20-24 G-Tracts, to N=12-24 G-Tracts. The parameters are the same as Figure 4: $K_{f,1}$ and $K_c$ are varied while $K_{f,2}$=2$K_{f,1}$, $K_f$ =60, $K_{f,N-1}$=80, and $K_{f,N}$=229. The colors in the phase diagram (top) correspond to the colors of the patterns of the S-parameter (bottom). The main patterns for the range N=20-24 G-Tracts (indicated by orange (4), green (1), and red (8)) persist over the entire range N=12-24 G-tracts. At the interfaces between these main patterns, other patterns are observed which bridge the transition between these main patterns with gradual shifts. For example, interface between the patterns shown in orange (4) and green (1), has two patterns shown in purple (3) and yellow (2). However, purple (3) differs from orange (4) in only one of the G-Tracts (N=12 to 13 transition), and yellow (2) differs from purple (3) in another G-Tract (N=16 to 17 transition). Finally, the yellow (2) pattern differs from the green pattern (1) in only one G-Tract (N=20 to 21 transition). Therefore, the transition from orange to green pattern occurs gradually through these small shifts in the pattern. In other words, the pattern shown in purple (3) consists of the pattern shown in orange (4) for N=13-20 and the pattern shown in green (1) for N=21-24. Similarly, the pattern shown in yellow (2) consists of the pattern shown in green (1) for N=12-20 and the pattern for orange (4) for N=20-24. The patterns shown in blue (5) and grey (9) show a similar interpolation at the interface between the patterns shown in green (1) and red (8). For the pattern in blue (5), there is cross-over from the pattern in orange (4) for N=12-20 to a different one for N=20-24. (Note this pattern becomes weaker at longer N as shown in Figure S2 (H).) This switching between one pattern at small N to another at larger N occurs at the interfaces between the patterns in orange (4), green (1), and red (8).

#### Entropy Calculations

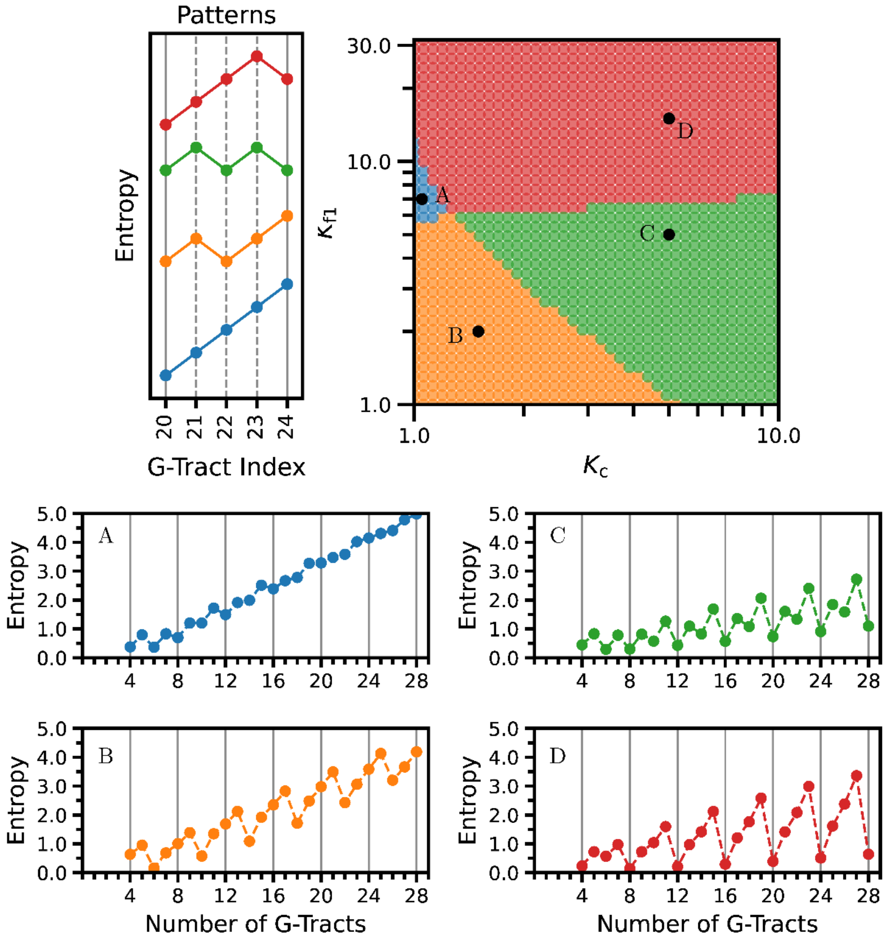


**Figure S4.** Computational patterns observed for the Entropy, $S/k_B = -\sum_\alpha p_\alpha \ln p_\alpha$, within the window of N=20-24 G-Tracts for varying $K_{f,1}$ and $K_c$ parameters. The parameters are the same as Figure 4: $K_{f,1}$ and $K_c$ are varied while $K_{f,2}$=2$K_{f,1}$, $K_f$ =60, $K_{f,N-1}$=80, and $K_{f,N}$=229. The top left panel shows the four patterns observed depending on whether the entropy increases or decreases as the length of the overhang is increased by one G-Tract. The phase diagram in top right shows the parameter range where different patterns are observed in colors that match the patterns on the left panel. Notice the phases overlap with those determined by the S-parameter shown in Figure 4, but the interfaces between these patterns are not the same. Still, the patterns in entropy illustrate that the frustration suggested by the S-parameter are reflected in the thermodynamics of the system. The Entropy as a function of the number of G-tracts is shown in parts A – D. The parameters for the plots in A-D are labeled on the phase diagram. A: $K_c$ = 0.105, $K_{f,1}$ = 7.0; B: $K_c$ = 15, $K_{f,1}$ = 2.0; C: $K_c$ = 5.0, $K_{f,1}$ = 5.0; D: $K_c$ = 5.0, $K_{f,1}$ = 15.0. The patterns in A and D are easy to interpret: In A, entropy increases with N. This is expected under conditions which the GQs are not very stabilized, ordered GQs have more initiation sites available as N increases. In D, the entropy has a minimum at N=[4n] and increases with additional G-Tracts until the next N=[4n + 3] as expected for very stabilized GQs with uniform stabilities at the ends. The patterns in B with local minima at N=[4n+2], and C with local minima N = [4n] and N=[4n+2] are harder to interpret. These are likely a result of the balance between formation of GQs at the 5' end which has lower stability and the rest of the telomere, similar to the explanation of the S-parameter patterns in shown in Figure 5.

#### FRET Melting Assay

We used FRET melting assay to determine the thermal stability of the duplex formed between the Cy5-PNA strand and a telomeric overhang that has a single binding site. A partial duplex DNA construct with the same duplex stem as those in FRET-PAINT studies (with a Cy3 at the same junction position) and overhang with sequence TTAGGGTTAG was used for these studies. This would effectively be a 1G-Tract construct in the nomenclature of our study (see Table S2 for sequence). The measurements were performed at the same ionic strength (150 mM KCl and 2 mM $MgCl_2$) and Cy5-PNA concentration (40 nM) as those used in FRET-PAINT assay.

Figure S5-A shows normalized FRET distributions, where the donor emission peaks are scaled to 1.0. An excitation wavelength of $\lambda$=532 nm was used while emission in 550-702 nm range was collected. The corresponding acceptor intensity, which is due to Cy5-PNA strand binding to the available G-Tract, decreases as the temperature increases, and reduces to background levels above ~30 °C. Figure S5-B shows acceptor peak as a function of temperature. Based on a sigmoidal fit, the thermal melting temperature is determined to be $T_m$=12.7±0.3 °C. These measurements are consistent with an empirical model that estimates the thermal melting temperature of PNA-DNA hybrid duplexes based on thermal melting temperatures of DNA-DNA duplexes. Based on this model, where oligo length, pyrimidine fraction, and DNA-DNA $T_m$ of our construct are used, we obtain $T_m$=13.7 °C. The estimates of this model typically deviated from experimental melting temperatures by a few °C.

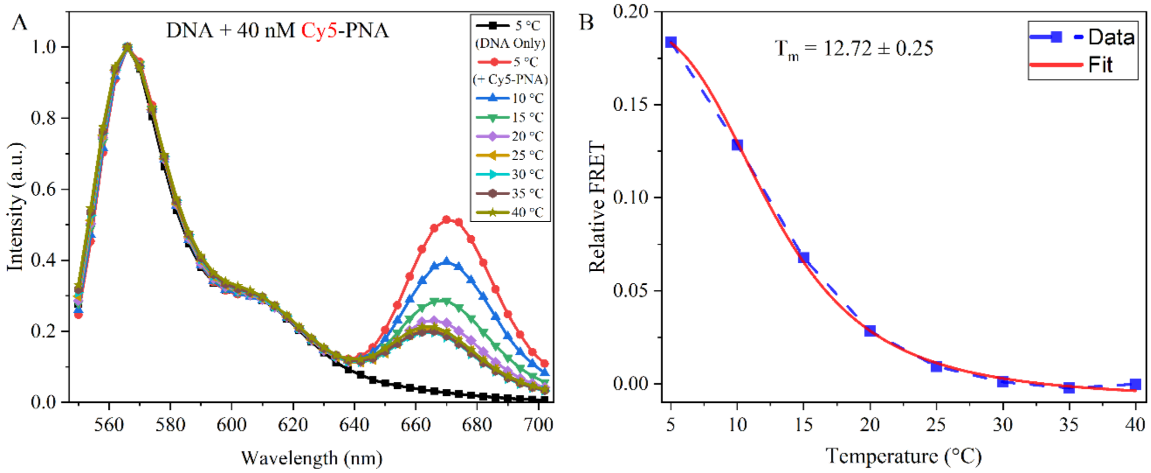


**Figure S5:** FRET melting measurements on 1G-Tract construct and Cy5-PNA. (A) Normalized FRET spectra at different temperatures. (B) Amplitude of acceptor emission peak as a function of temperature. A sigmoidal fit to the data results in $T_m$=12.7±0.3 °C.

#### Accessibility of 4G-Tract Construct Compared to Unfolded G-Tract

To get a feel for the level of accessibility we observe for telomeric overhangs, we performed comparative measurements on the 4G-Tract and 1G-Tract (same as that used for FRET melting assay) constructs. For these two constructs, we quantified binding frequencies (total number of binding events/total observation time per μM of Cy5-PNA strand), as shown in Figure S6. The binding frequency for the 4G-Tract construct is an order of magnitude lower compared to 1G-Tract construct. These data demonstrate that the GQ in 4G-Tract construct is folded in large majority of the constructs and remains folded over the 2-3 minute observation time. Unfolding of GQ would expose four bindings sites (four G-Tracts), while the reference construct (1G-Tract) contains a single binding site, further emphasizing the lower accessibility of 4G-Tract construct. These measurements also demonstrate that the Cy5-PNA probe does not present a significant disturbance to the GQ, in agreement with the implications of measurements of Figure S5.

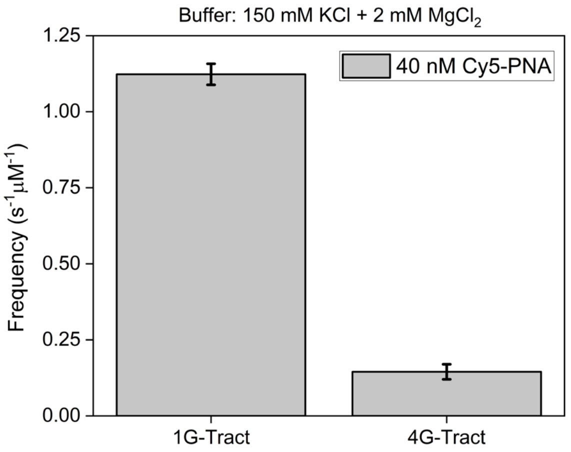


**Figure S6:** Comparative binding frequencies for 1G-Tract and 4G-Tract constructs per micromolar concentration of PNA (40 nM PNA was used in these measurements). The binding frequencies for the 4G-Tract construct are an order of magnitude smaller than those observed for the 1G-Tract construct, suggesting the GQ remains folded in vast majority of constructs over the 2-3 min observation time. These measurements also illustrate that any destabilization on GQ due to Cy5-PNA is relatively minor.

**Detection Sensitivity**

An important issue about the experimental approach is whether binding events to the vicinity of 3′-end (furthest from the donor fluorophore) are still within the detectable range for longer telomeric overhangs. To investigate this issue, we analyzed the shift in the accessibility patterns of Figure 2 to lower FRET efficiencies for [4n]G-Tract construct, i.e., 4, 8, 12, 16, 20, 24 and 28 G-Tract constructs. These constructs were chosen since the binding events are distributed throughout their overhang, rather than being concentrated in the vicinity of ssDNA/dsDNA junction where the majority of binding events result in a high FRET signal. Figure S7 shows the ratio of cumulative populations for $E_{FRET}<0.50$ and that for $E_{FRET}>0.50$. If detection sensitivity were lost beyond a certain overhang length, we would expect this ratio to remain constant beyond that length. However, that is not what we observed. The fraction of binding events at $E_{FRET}<0.50$ continues to increase. The increase is more significant at lower lengths and more gradual at longer lengths. This is expected due to two reasons: (i) the fractional change in the length of the overhang between consecutive constructs is smaller at longer lengths; (ii) large variations in donor-acceptor separation result in smaller variations in FRET efficiency at lower FRET range compared to mid-high FRET range.

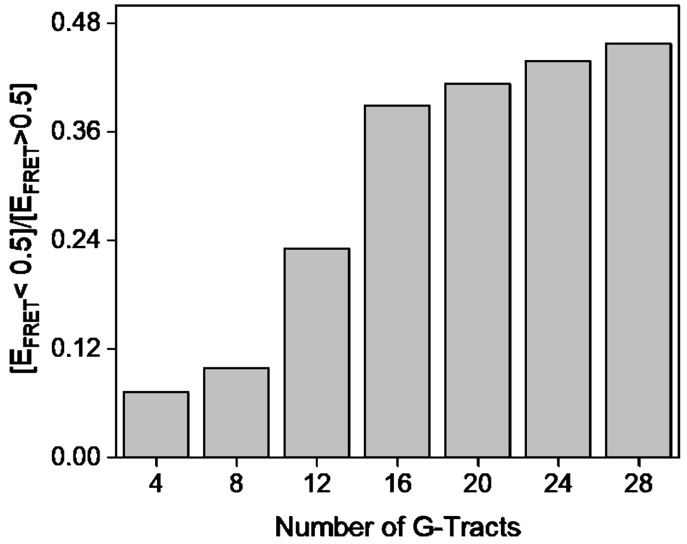


**Figure S7:** Ratio of cumulative FRET populations for $E_{FRET}<0.5$ and $E_{FRET}>0.5$. The continuous increase in this ratio with overhang length for the entire length range suggests our approach has the sensitivity to detect binding events even for the longer constructs.

#### Gel Purification

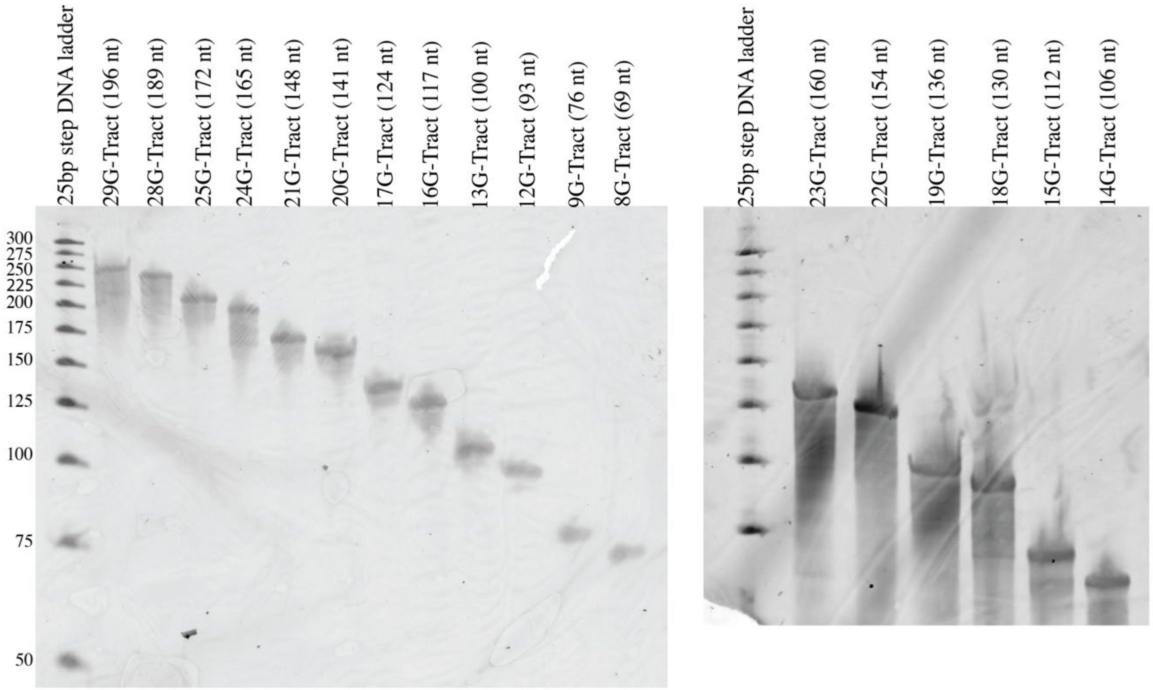


**Figure S8:** Denaturing polyacrylamide gel of purified DNA oligonucleotides. DNA oligos were purified via denaturing 10% polyacrylamide gel electrophoresis (PAGE). Full-length products were visualized by UV shadowing and the bands were excised from the gel. The DNA oligos were gathered via the crush and soak method by incubating and rocking of the gel slice overnight at room temperature in a solution of 300 mM NaCl, 10 mM Tris-HCl, and 0.1 mM EDTA (pH 7.4). The eluants were concentrated with sec-butanol and precipitated with 3 volumes of 100% ethanol. Salt was removed by washing twice, with 70% ethanol. After vacuum drying, the DNA pellet was dissolved in nuclease free water. Purified DNA oligos were resolved via denaturing 10% PAGE and gel was stained with SYBR gold for 15 minutes. Then the gel was visualized on a Typhoon FLA 9500 fluorescence imager (GE Life Sciences) by selecting Cy3 scanning mode.

**DNA Sequences**

The sequence of the Cy5-PNA probe was as follows: TAACCCTT-Cy5, the red nucleotides being complementary to the telomeric repeats. The partial duplex DNA (pdDNA) constructs were created by annealing a stem strand (18, 24 or 30 nt long) with a long strand that includes a complementary sequence to the stem strand and the telomeric overhang. The following strands were used to create the pdDNA constructs.

**Table S1:** Sequences for constructs used in measurements of Figure 1D and Figure 1F. The binding sites for Cy5-PNA are shown with red fonts.

| pd-DNA | Long Strand (Sequence in 5′–3′) | Stem |
|---|---|---|
| BS1-1GQ-BS2 | TGGCGACGGCAGCGAGGC TTA GGG TTA $(GGGT)_4$ TAGGGTTAG | Stem-18 |
| BS1-2GQ-BS2 | TGGCGACGGCAGCGAGGC TTA GGG TTA $(GGGT)_4$ TTA $(GGGT)_4$ TAGGGTTAG | Stem-18 |
| BS1-3GQ-BS2 | TGGCGACGGCAGCGAGGC TTA GGG TTA $(GGGT)_4$ TTA $(GGGT)_4$ TTA $(GGGT)_4$ TAGGGTTAG | Stem-18 |
| BS1-4GQ-BS2 | TTA GGG TTA $(GGGT)_4$ TTA $(GGGT)_4$ TTA $(GGGT)_4$ TTA $(GGGT)_4$ TAGGGTTAG | Stem-18 |
| 1GQ-BS | TGGCGACGGCAGCGAGGC TTA $(GGGT)_4$ TAGGGTTAG | Stem-18 |
| 2GQ-BS | TGGCGACGGCAGCGAGGC TTA $(GGGT)_4$ TA $(GGGT)_4$ TAGGGTTAG | Stem-18 |
| 3GQ-BS | TGGCGACGGCAGCGAGGC TTA $(GGGT)_4$ TA $(GGGT)_4$ TA $(GGGT)_4$ TAGGGTTAG | Stem-18 |
| 4GQ-BS | TGGCGACGGCAGCGAGGC TTA $(GGGT)_4$ TA $(GGGT)_4$ TA $(GGGT)_4$ TA $(GGGT)_4$ TAGGGTTAG | Stem-18 |
| 5GQ-BS | TGGCGACGGCAGCGAGGC TTA $(GGGT)_4$ TA $(GGGT)_4$ TA $(GGGT)_4$ TA $(GGGT)_4$ TA $(GGGT)_4$ TAGGGTTAG | Stem-18 |

**Table S2:** Sequences used for creating telomeric pdDNA constructs. The nucleotides in green fonts constitute the overhang. The subscripts designate the number of repeats, i.e. (GGGTTA)$_{4}$ refers to GGGTTAGGGTTAGGGTTAGGGTTA.

| pd-DNA | Long Strand (Sequence in 5′–3′) | Stem |
|---|---|---|
| 1G-Tract | TGGCGACGGCAGCGAGGCTTA(GGGTTA)$_{1}$ G | Stem-18 |
| 4G-Tract | TGGCGACGGCAGCGAGGCTTA(GGGTTA)$_{4}$ G | Stem-18 |
| 5G-Tract | TGGCGACGGCAGCGAGGCTTA(GGGTTA)$_{5}$ G | Stem-18 |
| 6-GTract | TGGCGACGGCAGCGAGGCTTA(GGGTTA)$_{8}$ G | Stem-30 |
| 7-GTract | TGGCGACGGCAGCGAGGCTTA(GGGTTA)$_{7}$ G | Stem-18 |
| 8G-Tract | TGGCGACGGCAGCGAGGCTTA(GGGTTA)$_{8}$ G | Stem-18 |
| 9G-Tract | TGGCGACGGCAGCGAGGCTTA(GGGTTA)$_{9}$ G | Stem-18 |
| 10G-Tract | TGGCGACGGCAGCGAGGCTTA(GGGTTA)$_{12}$ G | Stem-30 |
| 11-GTract | TGGCGACGGCAGCGAGGCTTA(GGGTTA)$_{13}$ G | Stem-30 |
| 12G-Tract | TGGCGACGGCAGCGAGGCTTA(GGGTTA)$_{14}$ G | Stem-30 |
| 13G-Tract | TGGCGACGGCAGCGAGGCTTA(GGGTTA)$_{15}$ G | Stem-30 |
| 14-GTract | TGGCGACGGCAGCGAGGCTTA(GGGTTA)$_{16}$ G | Stem-30 |
| 15G-Tract | TGGCGACGGCAGCGAGGCTTA(GGGTTA)$_{17}$ G | Stem-30 |
| 16G-Tract | TGGCGACGGCAGCGAGGCTTA(GGGTTA)$_{18}$ G | Stem-30 |
| 17G-Tract | TGGCGACGGCAGCGAGGCTTA(GGGTTA)$_{19}$ G | Stem-30 |
| 18G-Tract | TGGCGACGGCAGCGAGGCTTA(GGGTTA)$_{20}$ G | Stem-30 |
| 19G-Tract | TGGCGACGGCAGCGAGGCTTA(GGGTTA)$_{21}$ G | Stem-30 |
| 20G-Tract | TGGCGACGGCAGCGAGGCTTA(GGGTTA)$_{22}$ G | Stem-30 |
| 21G-Tract | TGGCGACGGCAGCGAGGCTTA(GGGTTA)$_{23}$ G | Stem-30 |
| 22G-Tract | TGGCGACGGCAGCGAGGCTTA(GGGTTA)$_{24}$ G | Stem-30 |
| 23G-Tract | TGGCGACGGCAGCGAGGCTTA(GGGTTA)$_{25}$ G | Stem-30 |
| 24G-Tract | TGGCGACGGCAGCGAGGCTTA(GGGTTA)$_{26}$ G | Stem-30 |
| 25G-Tract | TGGCGACGGCAGCGAGGCTTA(GGGTTA)$_{27}$ G | Stem-30 |
| 26G-Tract | TGGCGACGGCAGCGAGGCTTA(GGGTTA)$_{27}$ G | Stem-24 |
| 27G-Tract | TGGCGACGGCAGCGAGGCTTA(GGGTTA)$_{29}$ G | Stem-30 |
| 28G-Tract | TGGCGACGGCAGCGAGGCTTA(GGGTTA)$_{29}$ G | Stem-24 |

**Table S3:** Stem strands of 18, 24, or 30 nt length were used. After hybridizing with the long strand, the stem strand takes part in forming duplex DNA that is attached to the surface via biotin-streptavidin binding. All three strands have the same 18 nt common sequence (blue fonts). The 24-nt stem sequence has one additional C-Tract (CCCTAA, red fonts) while the 30-nt stem sequence has two additional C-Tracts (CCCTAACCCTAA, red fonts). These C-Tracts enable creating overhangs with different lengths.

| | |
|---|---|
| Stem-18 | Cy3-GCCTCGCTGCCGTCGCCA-biotin |
| Stem-24 | Cy3-CCCTAA GCCTCGCTGCCGTCGCCA-biotin |
| Stem-30 | Cy3-CCCTAACCCTAA GCCTCGCTGCCGTCGCCA-biotin |

**Table S4:** The number of DNA molecules ($N_B$) that showed at least one PNA binding event, $N_T$ represent the total number of DNA molecules that passed single molecule screening test and $N_N$ represent the number of DNA molecules that do not show any binding. The histograms in Figure 2 were constructed from PNA binding events to these $N_B$ molecules.

| Construct | $N_B$ | $N_T$ | $N_N = N_T - N_B$ |
|---|---|---|---|
| 4G-Tract | 74 | 380 | 306 |
| 5G-Tract | 92 | 414 | 322 |
| 6-GTract | 148 | 340 | 192 |
| 7-GTract | 91 | 309 | 218 |
| 8G-Tract | 58 | 297 | 239 |
| 9G-Tract | 148 | 349 | 201 |
| 10G-Tract | 100 | 171 | 71 |
| 11-GTract | 104 | 227 | 123 |
| 12G-Tract | 150 | 495 | 345 |
| 13G-Tract | 143 | 267 | 124 |
| 14-GTract | 120 | 340 | 220 |
| 15G-Tract | 158 | 271 | 113 |
| 16G-Tract | 238 | 510 | 272 |
| 17G-Tract | 181 | 351 | 170 |
| 18G-Tract | 161 | 350 | 189 |
| 19G-Tract | 195 | 371 | 176 |
| 20G-Tract | 135 | 214 | 79 |
| 21G-Tract | 136 | 278 | 142 |
| 22G-Tract | 126 | 315 | 189 |
| 23G-Tract | 200 | 359 | 159 |
| 24G-Tract | 108 | 319 | 211 |
| 25G-Tract | 146 | 248 | 102 |
| 26G-Tract | 135 | 194 | 59 |
| 27G-Tract | 134 | 222 | 88 |
| 28G-Tract | 145 | 260 | 115 |

**Table S5:** Peak $E_{FRET}$ values representing binding of Cy5-PNA to BS1 and BS2 in BS1-nGQ-BS2 constructs (based on Gaussian fits in Figure 1D).

| **DNA Construct** | **Peak 1 (BS1)** | **Peak 2 (BS2)** |
|---|---|---|
| BS1-1GQ-BS2 | 0.94 ± 0.01 | 0.84 ± 0.02 |
| BS1-2GQ-BS2 | 0.94 ± 0.01 | 0.78 ± 0.01 |
| BS1-3GQ-BS2 | 0.96 ± 0.01 | 0.71 ± 0.01 |
| BS1-4GQ-BS2 | 0.96 ± 0.01 | 0.50 ± 0.01 |

**Table S6:** Peak $E_{FRET}$ values representing binding of Cy5-PNA to BS in nGQ-BS constructs (based on Gaussian fits in Figure 1F).

| DNA Construct | Peak Position (BS) |
|---|---|
| 1GQ-BS | 0.93 ± 0.01 |
| 2GQ-BS | 0.90 ± 0.01 |
| 3GQ-BS | 0.77 ± 0.01 |
| 4GQ-BS | 0.67 ± 0.01 |
| 5GQ-BS | 0.63 ± 0.01 |
